# Applying drift diffusion models to rat gambling task data reveals divergent cognitive mechanisms underlying risky choice

**DOI:** 10.64898/2026.08.11.744251

**Authors:** Claire A. Hales, Catharine A. Winstanley

**Affiliations:** Department of Psychology, Djavad Mowafaghian Centre for Brain Health, University of British Columbia, Vancouver, BC, Canada

## Abstract

The rat gambling task (rGT) has been widely used to investigate the neural mechanisms underlying risky choice and motor impulsivity. Here, rats sample between four options (P1-P4) that vary in the size and probability of reward and time-out penalties. The optimal strategy is to avoid risky options that may yield higher per-trial gains, but deliver longer and more frequent time-outs. Previous reports suggest pairing wins with salient audiovisual cues increases risky decision making, but behavioural variation is high, and it is unclear whether motor impulsivity is also affected. Here we leveraged rGT data from over 750 rats to characterize behavioural performance across sex and cue condition. We compared different methods of classifying rats as optimal or risk-preferring, using either a unitary decision score variable or specific P-choice preference, and applied drift diffusion modeling (DDM) to explore whether divergent cognitive mechanisms underlie risky decision making across subgroups. We confirmed that risky choice is higher on the cued rGT, partly due to a greater proportion of risk-preferring rats, but also because net optimal decision-makers chose the risky options more often. Risk-preferring rats made more impulsive, premature responses regardless of cue condition, as did males. Optimal decision-makers made more premature responses when cues were present, such that premature response rates were higher overall on the cued rGT. DDM and response latency data suggest divergent cognitive processes underpinning risky decisions across sex. Wider decision boundaries were associated with both highly optimal and highly risky choice patterns, indicating risky choices are made deliberatively by highly risk-preferring individuals. Similar results were obtained regardless of classification method.

## Introduction

Decision-making under uncertainty involves a complex interplay of cognitive control, reward valuation, and impulsivity. The rat gambling task (rGT; Zeeb et al., 2009), loosely based on the Iowa Gambling Task used in humans (Bechara et al. 1994), is widely used to investigate these processes. In the task, rats are required to choose among four options (P1–P4) that differ in expected value based on reward magnitude, probability of reinforcement, and punishment duration. Two of these options (P1 and P2) offer smaller but more reliable gains and shorter time-out penalties, leading to greater overall reward accumulation across a session. In contrast, P3 and P4 offer larger per-trial rewards but are less frequently reinforced and carry longer penalties, reducing overall earnings. Most rats eventually learn to favor the advantageous P1/P2 choices, but a consistent minority persist in selecting the riskier, suboptimal P3/P4 options—a pattern mirroring individual differences observed in the IGT whereby a small proportion prefer the risky, disadvantageous options (Bechara et al., 1994; Brevers et al., 2013).

A growing body of work suggests that manipulations of rGT structure can significantly alter animals’ risk preferences. In particular, the addition of sensory cues paired with reward delivery, commonly referred to as the cued version of the rGT, has been shown to increase the proportion of rats exhibiting risk-preferring behavior compared to the standard uncued version (e.g., Zeeb et al., 2009; Barrus & Winstanley, 2016). While group-level changes in rGT performance in the presence of cues are well documented, understanding why some individuals deviate from optimal decision-making, in both cued and uncued tasks, remains a challenge. Similarly, less is known about how such task manipulations interact with individual traits such as biological sex, despite robust evidence from human studies that males and females often differ in risk-taking and impulsivity (Byrnes et al., 1999; Cross et al., 2011; van den Bos et al., 2013). Findings in rodents are mixed, with some suggesting that males are more impulsive or risk-prone (Orsini et al., 2016), while others report minimal or context-dependent differences (Orsini & Setlow, 2017).

This is further hindered by commonly used analytic strategies: most rGT studies rely on aggregate measures such as total decision scores (an aggregate measure across all four individual P-choices) and use this to classify rats into optimal or risk-preferring decision makers (Rivalan et al., 2013; Adams et al., 2017; Ferland & Winstanley, 2017; Langdon et al. 2019; Hathaway et al., 2021; Vonder Haar et al., 2022; Hales et al., 2025). However, such summary metrics may obscure meaningful individual differences with respect to which specific P-choice rats prefer, particularly in heterogeneous populations. To capture this variability, some researchers have adopted classification strategies based on dominant choice preferences—i.e., identifying rats that predominantly select one of the four options (P1–P4) across trials (Hultman et al., 2022; Tjernström & Roman, 2022; Lindberg et al., 2025), or using cluster analysis to identify choice-based phenotypes (Vonder Haar et al., 2022).

To move beyond surface-level behavioral patterns, computational modeling approaches - such as the drift diffusion model (DDM) - have emerged as powerful tools for dissecting decision-making into latent cognitive components: the speed of evidence accumulation (drift rate), decision caution (boundary separation), pre-choice bias (starting point), and non-decision time (e.g., factors unrelated to the decision process, such as sensory or motor processing). Importantly, DDM parameters can capture meaningful individual differences in impulsivity, reward sensitivity and psychiatric symptomatology. Alterations in drift rate have been linked to differences in value-based evidence accumulation, including biased reward representations observed in clinical populations (Moustafa et al., 2015; Pedersen et al., 2021; Pitliya et al., 2022; Shen et al., 2024). Reduced boundary separation along with altered drift rates have been linked to impulsive responding (Johnson et al., 2017; Sharma & Khan, 2018), reflecting that difference facets of impulsivity, such as impulsive action or impulsive choice, may arise from multiple computational pathways. By applying these models to large datasets, it becomes possible to move beyond coarse group conclusions and instead characterize the cognitive dynamics underlying choice patterns at a finer-grained level.

In the present study, we leveraged a large dataset of over 750 rats - a dataset that affords exceptional statistical power and resolution - to systematically investigate the cognitive and behavioral architecture of risky decision-making on the rGT. This large-n approach enabled us to go beyond aggregate metrics and detect subtle patterns and subtypes that are typically lost in smaller samples. First, we confirmed and extended prior findings showing that the cued version of the task increases the proportion of risk-preferring rats. Second, we examined whether sex differences emerge in behavioral outputs and model-derived decision parameters. Third, we explored the utility of diffusion modeling in parsing cognitive components underlying performance. Finally, we examine a fine-grained stratification based on individual rats’ preferred options (P1–P4) to identify subgroups with distinct behavioral and cognitive profiles.

By integrating high-resolution behavioral data with computational modeling across a uniquely large and diverse sample, this study provides novel insights into the mechanisms driving individual variability in risk-taking. This work not only enhances our understanding of how external and internal factors influence decision-making in rodents, but also lays the groundwork for improving translational models of psychiatric and neurocognitive disorders characterized by maladaptive choice behavior.

## Methods

### Subjects

Subjects were male and female Long Evans rats, either purchased from a commercial vendor (Charles River Laboratories, St. Constant, QC, Canada), or bred in-house. Rats were pair-, trio-or (for females only) quad-housed in a climate-controlled colony room on a reverse 12-hour light-dark cycle (lights off 08.00; temperature 21°C). Rats were food restricted to 85% of their free feeding weight and maintained on 14 g / 9 g (male/female) of standard rat chow, plus the sugar pellets earned in the task (∼5 g per day). Water was available ad libitum. Behavioural training began at least one week following the start of food restriction. All housing conditions and testing procedures were in accordance with the guidelines of the Canadian Council on Animal Care, and all protocols were approved by the Animal Care Committee of the University of British Columbia.

### Behavioural Data

Behavioural data were collated from twenty-six batches of rats that underwent behavioural training between 2012 and 2023. All rats were experimentally naïve during task acquisition. Within these twenty-six batches, 386 rats were female, 408 were male, 338 were trained on the uncued version of the rGT, and 456 were trained on the cued version. Rats with missing session data from the final three stable acquisition sessions were excluded. This left a final count of 776 rats: 376 female rats and 400 male rats; 325 rats trained on the uncued task and 451 rats trained on the cued task. A full breakdown of rat numbers by task and sex can be found in Table 1.

**Table 1.**
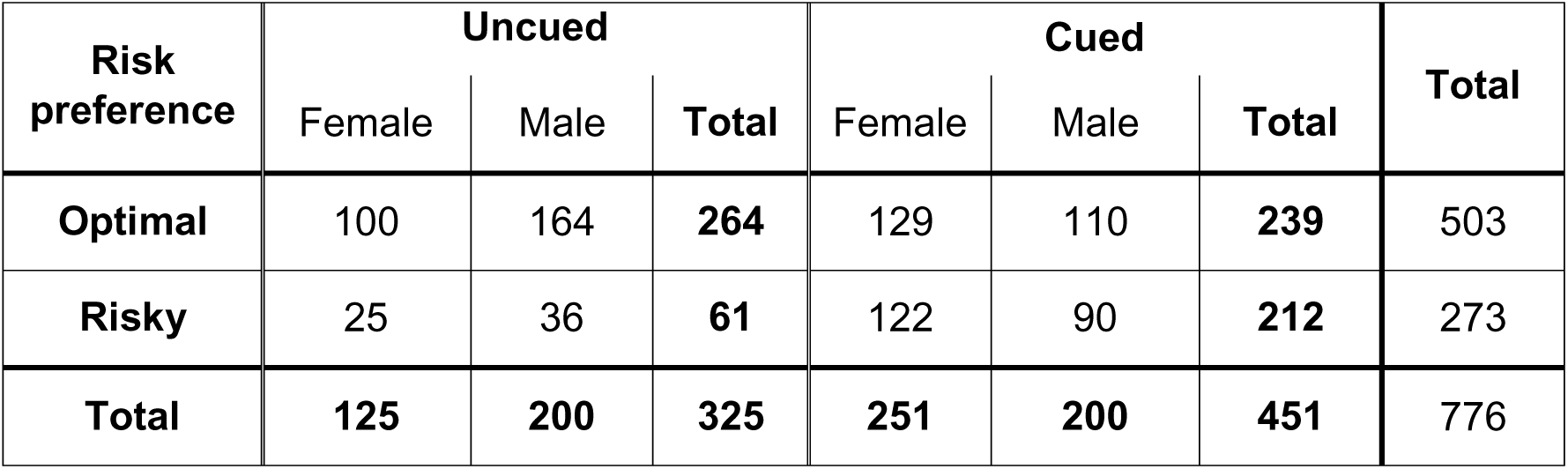
Breakdown of rats by risk preference. *Rats trained on the rGT were split into optimal or risky choice preference groups as per previous analyses. Numbers of optimal and risky rats are broken down by whether rats were trained on the uncued or cued rGT, and whether rats were male or female*.

### Rat Gambling Task – Apparatus and Training

Behavioural testing took place in 32 standard five-hole operant conditioning chambers, each enclosed within a ventilated sound-attenuating cabinet (Med Associates Inc., St. Albans, VT, USA), configured similarly to those previously described (Zeeb et al., 2009; Barrus & Winstanley, 2017). Operant boxes were controlled by software written in Med PC by CAW running on an IBM-compatible computer. Training sessions for each individual batch occurred at a consistent time within the dark phase of the light-dark cycle, although the time of training differed considerably within the dark phase between batches.

Task training was consistent across all batches. Rats first underwent two daily 30-minute habituation sessions in the operant chambers, during which sucrose pellets were present in the nose-poke apertures and food magazine. Initial behavioral training used a variant of the five-choice serial reaction time (5CSRT) task, in which rats learned to nose-poke in either holes 1, 2, 4, and 5 when the light inside was illuminated for 10 s. Correct responses resulted in delivery of one sugar pellet to the food magazine. Sessions were 30 minutes long and consisted of approximately 100 trials. Rats were trained until they reached a criterion of ≥ 50 correct responses with ≥ 80% accuracy and ≤ 20% omissions.

Rats then moved on to rGT training, which also consisted of 30-minute sessions. Figure 1A depicts the trial structure of the uncued and cued rGT (Barrus & Winstanley, 2017). Rats initiated a trial by making a nose-poke response within the illuminated food magazine. The tray light was extinguished, and a five-second inter-trial interval (ITI) followed, after which lights in apertures 1, 2, 4, and 5 in the five-hole array came on for a maximum of 10 s. A nose-poke response within one of the lit apertures within this time led to either reward or punishment with probabilities based on that aperture’s reinforcement schedule. Probability of reward varied among options (0.9, 0.8, 0.5, 0.4 for P1, P2, P3, P4), as did reward size (1, 2, 3, 4 sucrose pellets for P1, P2, P3, P4; Figure 1A). On unrewarded trials, a light flashed at 0.5 Hz within the chosen aperture, signalling a time-out penalty which varied in length depending on the aperture selected (5, 10, 30, 40 seconds for P1, P2, P3, P4; Figure 1A). Rats could earn maximal sucrose pellets by exclusively selecting the optimal P2 option, which offers a relatively high probability of a two-sucrose pellet reward (0.8) combined with short, infrequent time-out penalties (10 s, 0.2 probability). The P1 option is also relatively optimal, providing the next highest potential reward amount. Options P3 and P4 have higher per-trial gains of three or four sucrose pellets, but longer and more frequent time-out penalties which greatly reduces the occurrence of rewarded trials. Consistently selecting these options results in fewer sucrose pellets earned across the session and are therefore risker, disadvantageous options.

**Figure 1.**
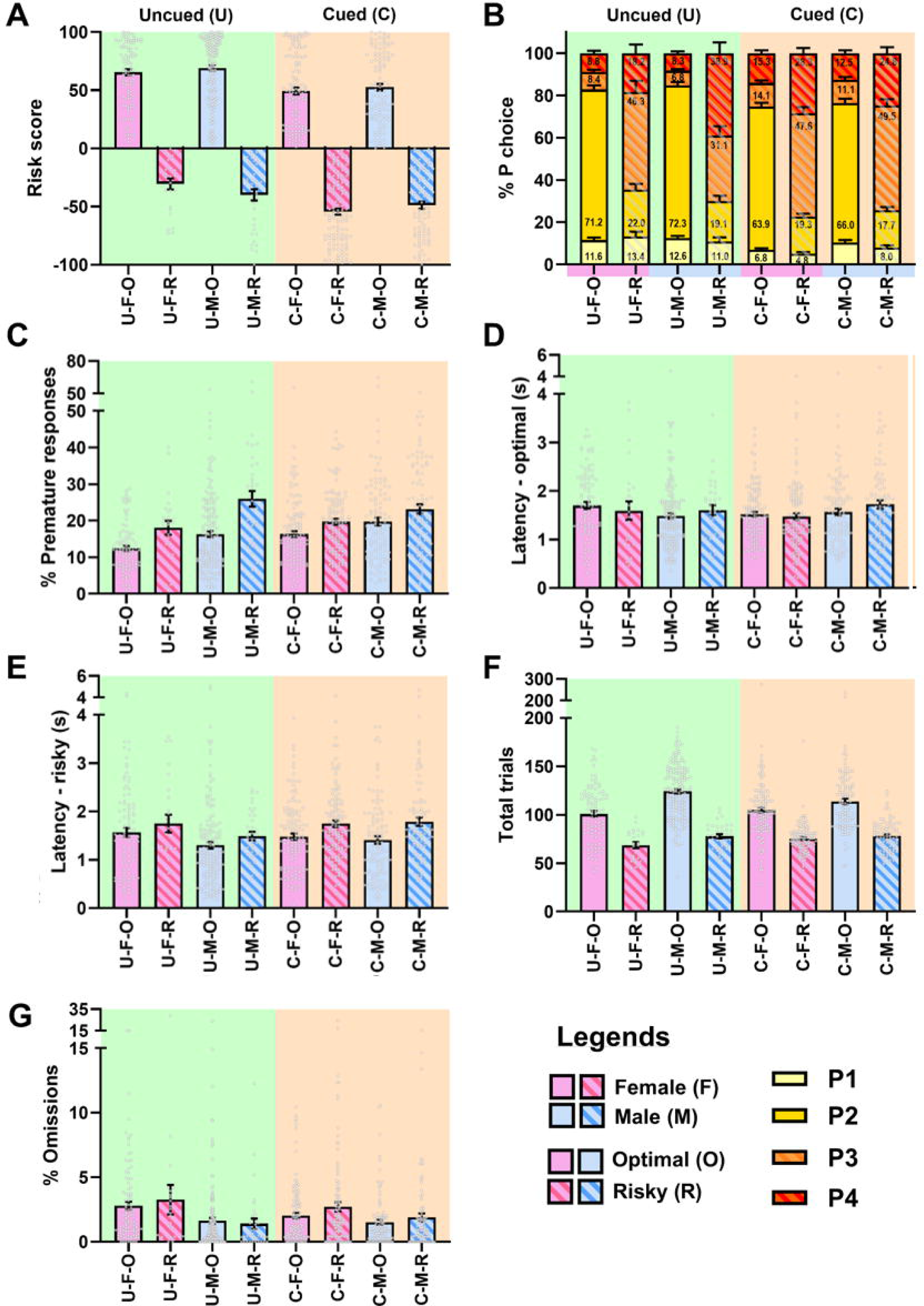
Behavioural performance on the cued and uncued rat gambling task classified by risk preference. Behavioural measures in rats performing the cued and uncued rat gambling task (rGT), grouped as optimal or risk-preferring decision-makers and separated by sex. (A) Decision scores were lower on the cued task compared with the uncued task. (B) Distribution of P-choices across the four options (P1–P4): Cues shifted choices towards riskier options, although the pattern differed according to sex and risk-preference classification. (C) Premature responses: Males made more premature responses than females, and risk-preferring rats were more impulsive than optimal decision-makers. Premature responding was elevated in optimal rats performing the cued task but was unaffected by cue condition in risk-preferring animals. (D) Latency to make optimal choices: Female rats made optimal choices more rapidly on the cued task, and were faster than males under cued conditions. (E) Latency to make risky choices: Males responded more rapidly than females, and optimal decision-makers selected risky options more quickly than risk-preferring rats. (F) Total trials completed: Males completed more trials than females, particularly on the uncued task, and optimal rats completed substantially more trials than risk-preferring rats. (G) Omissions: Females omitted more trials than males, whereas omission rates did not differ according to task variant or risk-preference classification. Data are presented as mean ± SEM, with individual data points overlaid. Significant effects are not displayed on this figure for clarity.

The cued rGT follows the same trial structure, except trials included a 2-second compound tone and light cue concurrent with reward delivery (Barrus & Winstanley, 2016). Cue complexity and variability scaled with reward size, such that the P1 cue consisted of a single tone and illuminated aperture, whereas the P4 cue featured multiple tones and flashing aperture lights presented in four different patterns across rewarded trials. For both tasks, the location of each option within the five-hole array was counterbalanced across rats to mitigate any potential side bias. Half of the rats in each batch were trained on version A (left to right arrangement: P1, P4, P2, P3) and the other half on version B (left to right arrangement: P4, P1, P3, P2). On both the cued and uncued rGT, if the rat did not respond in one of the lit apertures during 10 seconds, the trial was recorded as an omission, the food magazine was re-illuminated and the rat could initiate a new trial. A nose-poke response in the five-hole array before apertures were lit during the ITI was recorded as a premature response and punished by a five-second time-out period, during which the house light was illuminated and no trials could be initiated. At the end of the time-out, the food magazine was again illuminated, and rats could start another trial.

rGT training was split into forced-choice and free-choice sessions. After meeting criteria for 5CSRT training, rats completed 7 forced-choice sessions, during which only one of the four P1-P4 options were presented in trial. This ensured that rats had equal exposure to each reinforcement contingency prior to free-choice training, when all four options were available on each trial. All batches of rats received 5–6 training sessions per week, and continued on free-choice rGT training until stable performance was achieved i.e. when the percentage choice of P1 to P4 did not differ across three sessions as analyzed with repeated measures analysis of variance (ANOVA) with session as the within-subjects factor.

### Behavioural data analysis

Behavioral data from the three stable acquisition sessions at the end of rGT free choice training were analysed per session for each rat. As per previous methods (Zeeb et al., 2009; Barrus & Winstanley 2016; 2017), a decision score was calculated as follows: the percent of optimal (P1+P2) minus risky (P3+P4) choices. Rats with a positive decision score are classified as optimal, while rats with a negative decision score are classified as risk-preferring. Other behavioral measures were also calculated for each session for each rat, including: total trials completed, average latency to choose an optimal option (P1 and P2), average latency to choose a risky option (P3 and P4), percentage of omissions, percentage of premature responses, as well as percentage choices of individual options (P1-4).

### Cluster analysis

To investigate whether groupings rats into distinct optimal and risk-preferring groups based on decision score is a valid approach, unsupervised K-means clustering (using the kmeans function in Matlab) was carried out. Clustering was run on two different subsets of data. First, to check whether other behavioral or cognitive measures might be important for group classifications, clustering was run including decision score, optimal choice latency, risky choice latency and % premature responses, along with the four main drift diffusion model parameters: drift rate, starting point, boundary and non-decision time. This will be referred to as all-measures clustering. Second, to address claims in the literature that P-choice preference patterns may be important in distinguishing individual’s choice strategies, and to allow closer comparison with alternative previously used analysis methods (Hultman et al., 2022; Tjernström & Roman, 2022; Vonder Haar et al., 2022; Lindberg et al., 2025), we ran clustering analysis using only percentage choice for the four different P-choice options. This will be referred to as P-choice clustering. For both approaches, data were averaged for the three stable acquisition sessions to reduce the effect of individual variation across sessions within the clustering approach. The optimal number of clusters was determined using all four methods within the evalclusters function in Matlab, (Calinski-Harabasz criterion, Davies-Bouldin criterion, Gap Value criterion and Silhouette criterion). For the all-measures clustering, two different optimal cluster numbers were found, with consensus between the Calinski-Harabasz / Gap Value methods (k=5) and Davies-Bouldin / Silhouette methods (k=2; Table S1). Splitting the data using the clusters found with k=2 resulted in rats essentially being classified using traditional optimal/risk-preferring groupings, whilst the k=5 cluster classification stratified rats into more specific clusters along the same axis (i.e. optimal to risk-preferring), resulting in groups that corresponded to “high-optimal”, “mid-optimal”, low-optimal”, “low-risky” and “high-risky” classifications. Given that these results were largely consistent with a categorization based on positive or negative decision score, the results of the all-measures analysis are shown in Supplementary Information. The P-choice clustering resulted in k=3 as the optimal number of clusters (consistent across all evalclusters methods). To evaluate what pattern of P-choice preference these three groups represented, we manually classified rats based on their single top P-choice preference. The numbers of rats within each manually classified P-choice preference group (P1, P2, P3 or P4 preferring) were then compared for each of the three cluster groups, and cluster groups named based on the majority single P-choice preference (Pchoice2, Pchoice3, Pchoice4) in each.

### Drift diffusion modeling

The diffusion model was fit to behavioural data from the choice rGT using fast-dm-30.2 (Voss & Voss, 2007; 2008; Voss et al., 2010; 2015). P1 and P2 choices were combined to characterise an optimal choice, and P3 and P4 responses were combined to characterise a risky choice. Optimal/risky choice, and response times (RTs) for these choices were input to the model. Figure 1A shows the task structure, with the behavioral measures used for modeling highlighted. Data from individual rats and sessions (three stable acquisition sessions) were modeled separately. Fast-dm calculates predictive cumulative distribution functions (CDFs) for choices and RTs, and then uses a partial differentiation equation solver to model the evolution of the probability distribution forward in time. Parameters are optimised by using an implementation of the Nelder-Mead method (Nelder & Mead, 1965). Further details about diffusion modelling using fast-dm can be found in Voss et al. (2015). As carried out previously (Hales et al., 2016; 2017; 2024; 2025), and following recommendations given in Voss et al. (2015), model fit was assessed using Kolmogorov-Smirnov (KS) test statistics output by fast-dm-30.2. The KS test statistic is the maximum absolute vertical distance between the empirical and the predicted CDFs of the RT distributions. For multiple trials in a task, n, it is computed as:

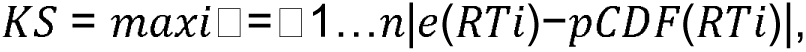

where *eCDF* and *pCDF* are the empirical and predicted CDFs, respectively, and *RTi* is the response latency in trial *i*. p-values < 0.05 indicate that the model does not demonstrate a good fit. Multiple models were tested using different combinations of parameters that were fit to all trials from a subset of the data (n = 100) in which an optimal or risky choice was made (i.e. premature and omission trials were excluded) to identify the parameter combination that produced best model fits. Validation of this best fitting model was carried out on the full behavioural dataset (n = 776) and used to model the behavioural data. Using the best fitting model, all sessions for all rats had model fit KS statistics > 0.132, and so no data were excluded from the analysis based on model fit.

The best fitting model had six parameters: starting point (zr), boundary separation (a), drift rate (v), non-decision RT (t0), the difference in speed of response execution between the two responses (d), and the variability in the starting point (szr). The other parameters that can be fit within fast-dm-30.2: variability in drift rate (sv), variability in non-decision RT (st0) and percentage of contaminants (p); were set to 0. In the model, the upper boundary represents an optimal choice, while the lower boundary represents a risky choice.

## Statistical analysis

Behavioral data was analyzed using custom code written in MATLAB. Behavioral measures and diffusion model parameters were averaged for individual rats across the three stable sessions for statistical analysis. Behavioral measures and diffusion model parameters also underwent two different group analyses – the first using the tradition optimal/risk-preferring groupings from decision score, and the second using the subgroups based on p-choice clustering. Pearson’s Chi-Square was used to test whether proportions of optimal and risk-preferring rats differed by task and sex. For the risk preference analysis, measures were analyzed using univariate ANOVAs with task (uncued or cued), sex (male or female), and risk preference (optimal or risk-preferring) as between-subjects factors. For percentage choices of P1-P4, a mixed ANOVA with the same between-subjects factor was used for analysis, but also included P-choice as the within-subjects factor. For the P-choice clustering analysis, measures were analyzed using univariate ANOVAs with task (uncued or cued), sex (male or female), and P-choice cluster subgroup as between-subjects factors. P-choice percentages did not undergo statistical analysis as the subgroups were split based on P-choice percentage criteria.

Given the large dataset, we also looked at whether there were any relationships between diffusion model parameters and the behavioural measures that have primarily been deemed the main measures of interest in previous work: decision score, to capture risky decision making, and percentage premature responses, to capture motor impulsivity. Linear relationships between model parameters and decision score / percentage premature responses were measured using Pearson’s Correlation Coefficients, with Pearson’s *r* and associated p-value reported for significant correlations. Quadratic relationship was measured using a second order polynomial with least squares fit, with *r^2^* value reported.

Post-hoc comparisons were performed with Bonferroni correction for multiple comparisons for significant two-way main effects or interactions (task, sex, risk preference), and Tukey’s HSD post-hoc comparison for cluster main effects or interactions. Huynh-Feldt corrections were used to adjust for violations of the sphericity assumption on rmANOVAs.

Statistics are reported with the ANOVA F-value (degrees of freedom, error) and p-value as well as any post-hoc p-values. Statistical tests on behavioral data were performed using SPSS 28.0.0.0 for Windows (IBM SPSS Statistics) and GraphPad Prism 10.4.0 for Windows (GraphPad Software, USA) was used for visualizations.

## Results

### Analysis using risk-preferring and optimal categories based on score (RP/OPT)

Within the uncued task, a similar proportion of females were optimal (80.0%) as males (82.0%; Pearson Chi-Square: χ_(1)_= 0.202, p = 0.653; Table 1). There was also no difference in proportion of optimal female (51.4%) or male (55.0%) rats within the cued task (Pearson Chi-Square: χ_(1)_ = 0.581, p = 0.446; Table 1). As expected from previous work, more rats were designated as risk-preferring on the cued task (47.0%) compared to the uncued task (18.8%; Pearson Chi-Square: χ_(1)_ = 66.045, p < 0.001; Table 1).

### Analysis using P-choice categories (PchoiceX)

When classified manually, the majority of rats (n = 490) were P2 preferring, whilst 17 were P1 preferring, 173 were P3 preferring, and 96 were P4 preferring. Table 2 describes how many rats in each of these subgroups were trained on the cued vs uncued task, and how many were male vs female. Comparing numbers of rats within each P-choice category in the three P-choice clustering subgroups, it became apparent that the three groups correspond to predominantly P2 preferring rats (408 out of 410); 2) P3 preferring (170 out of 202); and mostly P4 preferring (85 out of 164). We will therefore label these groups Pchoice2, Pchoice3, and Pchoice4. There were more Pchoice2 rats on the uncued task (57.1%), and fewer Pchoice3 (22.8%) and Pchoice4 (27.4%) compared to the cued task (Pearson Chi-Square: χ_(2)_ = 83.232, p < 0.001; Table 2). There were more male rats than expected in the Pchoice2 group (55.9%), and fewer male rats in the Pchoice3 group (44.6%; Pearson Chi-Square: χ_(2)_ = 7.305, p = 0.026; Table 2), but equivalent numbers of male and female rats in Pchoice 4 (49.4% vs 50.6%; Table 2).

**Table 2.** Breakdown of rats by P-choice preference. *Rats trained on the rGT were split into subgroups based on P-choice preference. Number of rats in each subgroup is broken down by whether rats would have been classified as optimal or risky using the traditional split of risk scores greater or smaller than 0 respectively; whether rats were trained on the uncued or cued rGT, and whether rats were male or female*.

| P-choice preference | Optimal | Risky | Uncued | Cued | Female | Male | Total |
| --- | --- | --- | --- | --- | --- | --- | --- |
| Pchoice2 | 410 | 0 | 234 | 176 | 181 | 229 | 410 |
| Pchoice3 | 33 | 169 | 46 | 156 | 112 | 90 | 202 |
| Pchoice4 | 60 | 104 | 45 | 119 | 83 | 81 | 164 |

### Decision score

#### RP/OPT

Decision scores were more negative in the cued task (task: F_1,768_ = 42.629, p < 0.001; Figure 1A), replicating previous work (e.g. Barrus & Winstanley, 2016; Betts et al., 2021; Chernoff et al., 2021). This was the case even within risk-preferring and optimal subgroups, with the exception of risk-preferring males (task*sex*risk preference: F_1,768_ = 4.981, p = 0.026; cued vs uncued for risk-preferring males: p = 0.123; optimal males: p < 0.001; risk-preferring females: p < 0.001; optimal females: p < 0.001; Figure 1A).

#### PchoiceX

Decision scores varied by P-choice category as expected (P-choice category: F_2,764_ = 722.478, p < 0.001; Figure 2A). Pchoice2 rats had the most positive decision scores (ps < 0.001), and Pchoice3 the most negative (ps < 0.001). Similar to the RP/OPT analysis, scores within each group were similar across sex. Again, across categories, decision scores were more negative on the cued task (task*P-choice category: F_2,764_ = 3.600, p = 0.028; P2: p < 0.001, P3: p < 0.001, P4: p = 0.051).

**Figure 2.**
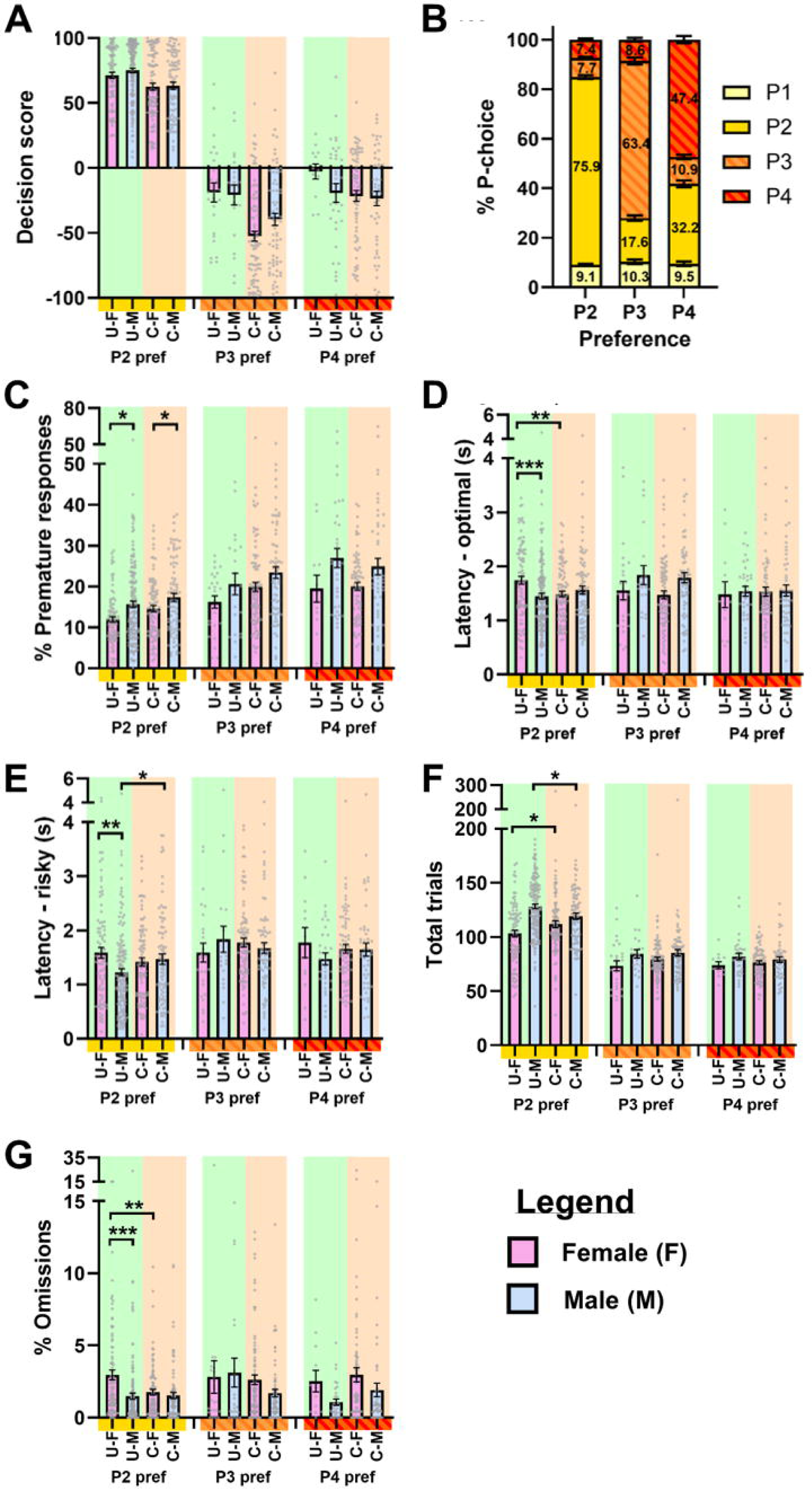
Behavioural performance on the cued and uncued rat gambling task classified by P-choice category. Behavioural measures in rats performing the cued and uncued rat gambling task (rGT), grouped according to choice category (Pchoice2, Pchoice3, or Pchoice4) and separated by sex. (A) Decision scores differed between P-choice categories, with Pchoice2 rats displaying the most advantageous decision-making and Pchoice3 rats the riskiest. Across categories, decision scores were generally lower on the cued task. (B) Distribution of P-choices across the four options (P1–P4): Cues altered P-choices within Pchoice2 and Pchoice3 groups but not Pchoice4 animals. Sex differences were only observed in the Pchoice4 group with females choosing P1 less frequently and P2 more frequently than males. (C) Premature responses: Pchoice2 rats made fewer premature responses than both Pchoice3 and Pchoice4 rats. (D) Latency to make optimal choices: Overall latencies did not differ between P-choice categories, although male Pchoice3 rats were slower to make optimal choices than males in the other groups. Within Pchoice2, females on the uncued task had longer response latencies. (E) Latency to make risky choices: Pchoice2 rats made risky choices more rapidly than Pchoice3 and Pchoice4 rats. (F) Total trials completed: Pchoice2 rats completed more trials than either Pchoice3 or Pchoice4 animals. Within the Pchoice2 group, females completed more trials on the cued task, whereas males completed more trials on the uncued task. (G) Omissions: More omissions were made by female rats in the Pchoice2 group on the uncued task. Data are presented as mean ± SEM, with individual data points overlaid. *p<0.05, **p<0.01, ***p<0.001.

As such, not only does the cued task increase the number of risk-preferring or Pchoice3 rats, but decision-making within subgroups is on average riskier in the presence of win-paired cues.

### P-choice

#### RP/OPT

Preference for P-choice differed across task, sex and risk preference (P-choice*task*sex*risk preference: F_2.335,1793.347_ = 5.520, p = 0.002; Figure 1B). We first compared performance of each group (male/female, optimal/risk-preferring) *between* cue conditions. Optimal females chose P1 less frequently (p < 0.001), with a corresponding increase in P4 (p = 0.036), if trained on the cued task. Risk-preferring females likewise chose P4 more frequently at the expense of P1 when wins were cued (ps < 0.015). Choice of P2 and P3 remained similar across task variant. In contrast, optimal males chose P2 less on the cued task (p = 0.005), and P3 and P4 more often (ps < 0.029), whereas risk-preferring males chose P3 more often (p < 0.001), and surprisingly P4 less often (p = 0.011).

We then compared male/female and risk-preferring/optimal groupings *within* each cue condition. When trained on the uncued task, risk-preferring females preferred P3 over P4 whereas the inverse was true for risk-preferring males (ps < 0.001). In contrast, optimal males and females looked very similar in their P-choice patterns (ps > 0.373). When trained on the cued task, females consistently prefer P1 less than males across both risk-preferring (p = 0.042) and optimal groupings (p = 0.006), but no other P-choice was significantly different between the sexes (ps > 0.205).

Zooming out further, and collapsing across sex, the proportions of P1 and P2 choices were both significantly lower in optimal rats performing the cued task (ps < 0.004), with correspondingly greater choice of P3 (p = 0.036) and P4 (p = 0.004). For risk-preferring animals, only choice of P1 and P3 differed, with fewer P1 choices and more P3 choices on the cued task (ps < 0.001, P2 and P4: all ps > 0.202).

#### PchoiceX

There were differences across task (P-choice*task*P-choice category: F_2.681,2048.662_ = 2.856, p = 0.012) and sex (P-choice*sex*P-choice category: F_2.681,2048.662_ = 2.423, p = 0.030) in the relative amount of different P-options chosen within PchoiceX groups. Comparing between cue conditions, Pchoice2 rats picked P1 less (p < 0.001), and P3 more (p = 0.021) on the cued task; Figure 2B). Pchoice3 rats showed the opposite pattern for numbers of P1 (p < 0.001) and P3 choices (p < 0.001), whilst also picking P2 less on the cued task (p = 0.029). There were no differences in Pchoice4 rats.

When breaking P-choices within PchoiceX group down by sex, there was only a difference in the Pchoice4 group, with females choosing P1 less (p = 0.037) and P2 more (p = 0.002) than males (Figure 2B).

### Motor impulsivity

As has been reported previously, male rats made more premature responses than females (sex: F_1,768_ = 29.203, p < 0.001; Figure 1C).

#### RP/OPT

Risk-preferring rats made more premature responses than optimal decision-makers (risk preference: F_1,768_ = 40.793, p < 0.001; Figure 1C). Given that more rats prefer the risky options on the cued rGT, it may seem unsurprising that premature response rates are higher on the cued task (F_1,768_ = 3.844, p = 0.050). However, this latter effect is driven by a difference in *optimal rats only* (task*risk preference interaction: F_1,768_ = 6.139, p = 0.013; cued vs uncued for optimal rats: p < 0.001; Figure 1C); risk-preferring animals make comparable amounts of premature responses regardless of whether cues are present (p = 0.686).

#### PchoiceX

Pchoice2 rats made fewer premature responses than Pchoice3 or Pchoice4 (P-choice category: F_2,764_ = 34.690, p < 0.001; Figure 2C ps < 0.001).

Both classification schemes therefore show that animals that make more advantageous decisions make fewer impulsive responses in general.

#### Response latencies

Female rats were quicker to make optimal choices on the cued task (task*sex: F_1,764_ = 4.584, p = 0.033, uncued vs cued: p = 0.044), and female rats were quicker to make optimal choices compared to males on the cued task only (p = 0.020; Figure 1D). For risky choice latencies, male rats were quicker to make risky choices across both tasks (sex: F_1,758_ = 2.573, p = 0.047; Figure 1E).

##### RP/OPT

Optimal choice latencies did not differ across groups, but optimal decision-makers made risky choices more quickly than risk-preferring rats across both task variants (risk preference: F_1,758_ = 13.353, p < 0.001; Figure 1D).

##### PchoiceX

Optimal choice latencies did not differ across subgroups (P-choice category: F_2,760_ = 1.589, p = 0.205; Figure 2D), however there was an interaction with sex (F_2,760_ = 4.791, p = 0.009). In female rats there were no differences in optimal choice latencies between subgroups (ps > 0.272), whereas in male rats, optimal choice latencies were longer in Pchoice3 rats (ps < 0.021). Within Pchoice2, females were slower to make optimal choices on the uncued task (task*sex: F_1,406_ = 7.812, p = 0.005), both compared to males on the same task (p < 0.001), and compared to their Pchoice2 counterparts on the cued task (p = 0.008).

Pchoice2 rats were quicker to make risky choices than Pchoice3 and Pchoice4 animals (P-choice category: F_2,754_ = 8.092, p < 0.001: Pchoice2 vs PChoice3: p < 0.001; Pchoice2 vs PChoice4: p = 0.006 Figure 2E). Within Pchoice2, female rats were slower to make risky choices on the uncued task compared to males (task*sex: F_1,396_ = 6.058, p = 0.014; uncued males vs uncued females: p = 0.001), and males were quicker to make risky choices on the uncued task compared with cued (p = 0.034).

Across both classification schemes, more advantageous decision making was therefore associated with quicker risky choices. The observation that female rats were generally slower to make a choice appears to be driven by Pchoice 2 animals.

#### Task engagement: trials completed and omitted

Male rats completed more trials than female rats (sex: F_1,768_ = 16.823, p < 0.001), and this difference was more pronounced in the uncued task (task*sex: F_1,768_ = 5.711, p = 0.017; Figure 1F). Male rats also made fewer omissions than females (sex: F_1,768_ = 18.825, p < 0.001; Figure 1G), but there were no differences by task, risk preference, or P-choice category.

##### RP/OPT

Optimal decision-makers completed more trials than risk-preferring rats (risk preference: F_1,768_ = 275.691, p < 0.001). Within optimal rats only, male rats completed more trials than females (sex*risk preference: F_1,768_ = 5.739, p = 0.017; optimal males vs females: p < 0.001).

##### PchoiceX

Pchoice2 rats completed more trials than other subgroups (P-choice category: F_2,764_ = 170.945, p < 0.001; pairwise comparisons: ps < 0.001; Figure 2F). Within this group, females completed more trials on the cued task whereas males completed more trials on the uncued task (task*sex: F_1,406_ = 10.029, p = 0.002; female cued vs uncued: p = 0.034; male cued vs uncued: p = 0.018).

As such, in both classification systems, advantageous decision making was associated with more trials completed, which is as expected given that fewer, shorter time-out penalties would be experienced.

#### Do diffusion model parameters relate to level of risky choice or motor impulsivity?

Correlations were used to test whether diffusion model parameters were related to decision scores or percentage of premature responses. Decision score was significantly positively correlated with both drift rate (Pearson’s *r* = 0.848, p < 0.001; Figure 3A) and starting point (Pearson’s *r* = 0.442, p < 0.001; Figure 3B), indicating that more positive drift rates and more positive starting points (towards optimal options) were associated with more optimal decision scores. There was a U-shaped relationship between the boundary parameter and decision score (quadratic least squares *r^2^* = 0.316; Figure 3C), with wider decision boundaries being associated with both highly optimal and highly risky decision scores. There was no relationship between decision score and non-decision time (Figure 3D).

**Figure 3.**
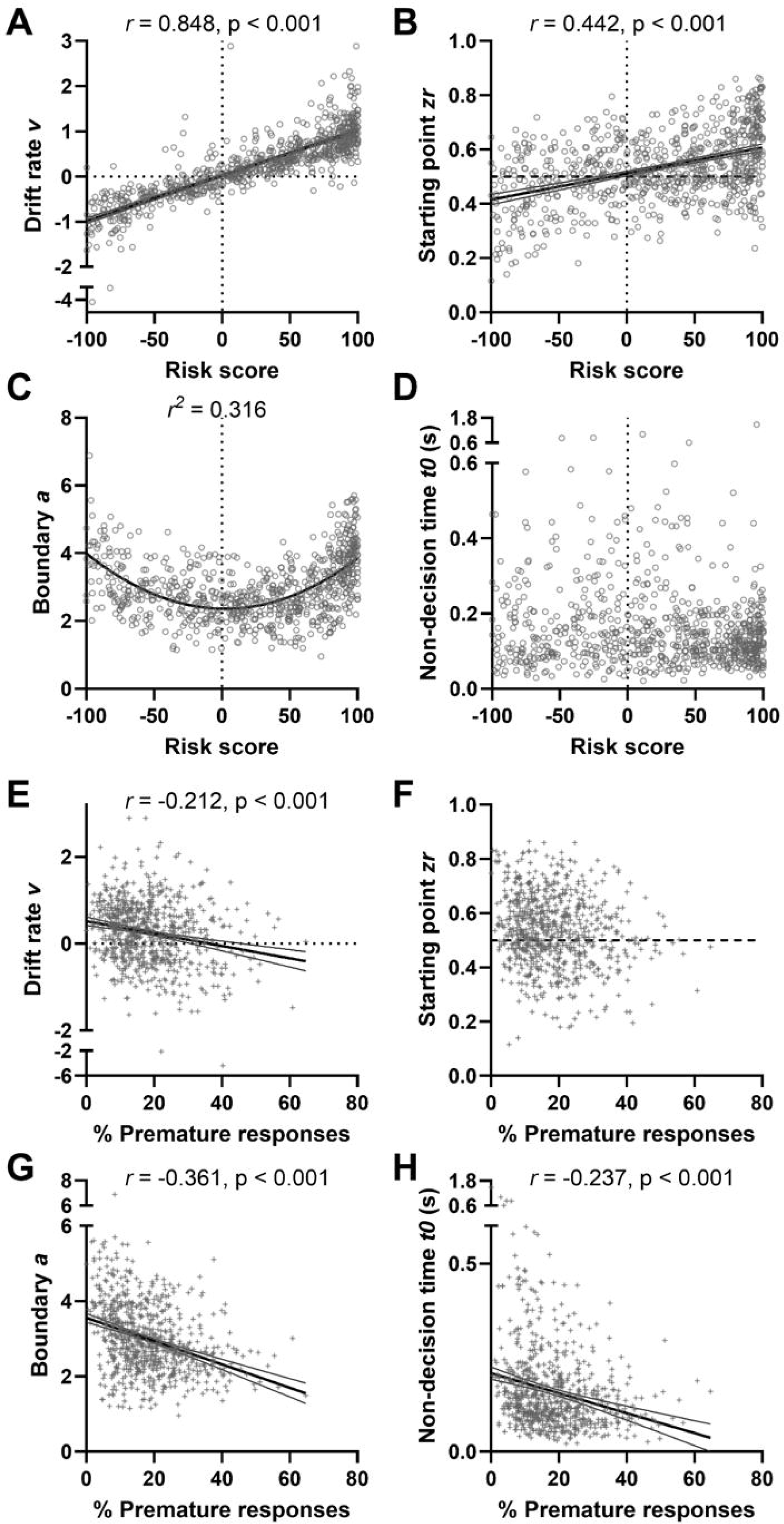
Relationship between diffusion model parameters, decision-making and impulsivity. Correlations between diffusion model parameters and behavioural measures on the rat gambling task (rGT). (A) Drift rate was positively correlated with decision score, indicating that more positive evidence accumulation toward advantageous options was associated with more optimal decision-making. (B) Starting point was also positively correlated with decision score, demonstrating that an initial bias toward advantageous options predicted more optimal performance. (C) Boundary separation showed a U-shaped relationship with decision score, such that wider decision boundaries were associated with both highly optimal and highly risky choices. (D) No relationship was observed between non-decision time and decision score. (E) Premature response rates were negatively correlated with drift rate, indicating that greater motor impulsivity was associated with evidence accumulation biased toward risky choices. (F) There was no relationship between premature responding and starting point. (G) Premature responding was negatively correlated with boundary separation, suggesting that more impulsive rats had narrower decision thresholds. (H) Premature responding was also negatively correlated with non-decision time, indicating that rats with higher levels of motor impulsivity had shorter non-decision processes. Data points represent individual rats; lines indicate the best-fitting linear relationship (A, B, E, G, H) or quadratic fit (C).

Premature response rates were negatively correlated with drift rate (Pearson’s *r* = - 0.212, p < 0.001; Figure 3E), boundary (Pearson’s *r* = -0.361, p < 0.001; Figure 3G) and non-decision time (Pearson’s *r* = -0.237, p < 0.001; Figure 3H). As such, increased premature responding is associated with more negative drift rates (towards risky choices), narrower decision boundaries and shorter non-decision times. There was no relationship between premature responding and decision starting points (Figure 3F).

### Diffusion model parameters analyzed by rat subgroups

#### Drift rate

Overall, drift rates were more negative in females compared to males (sex: F_1,768_ = 4.729, p = 0.030; Figure 4A).

**Figure 4.**
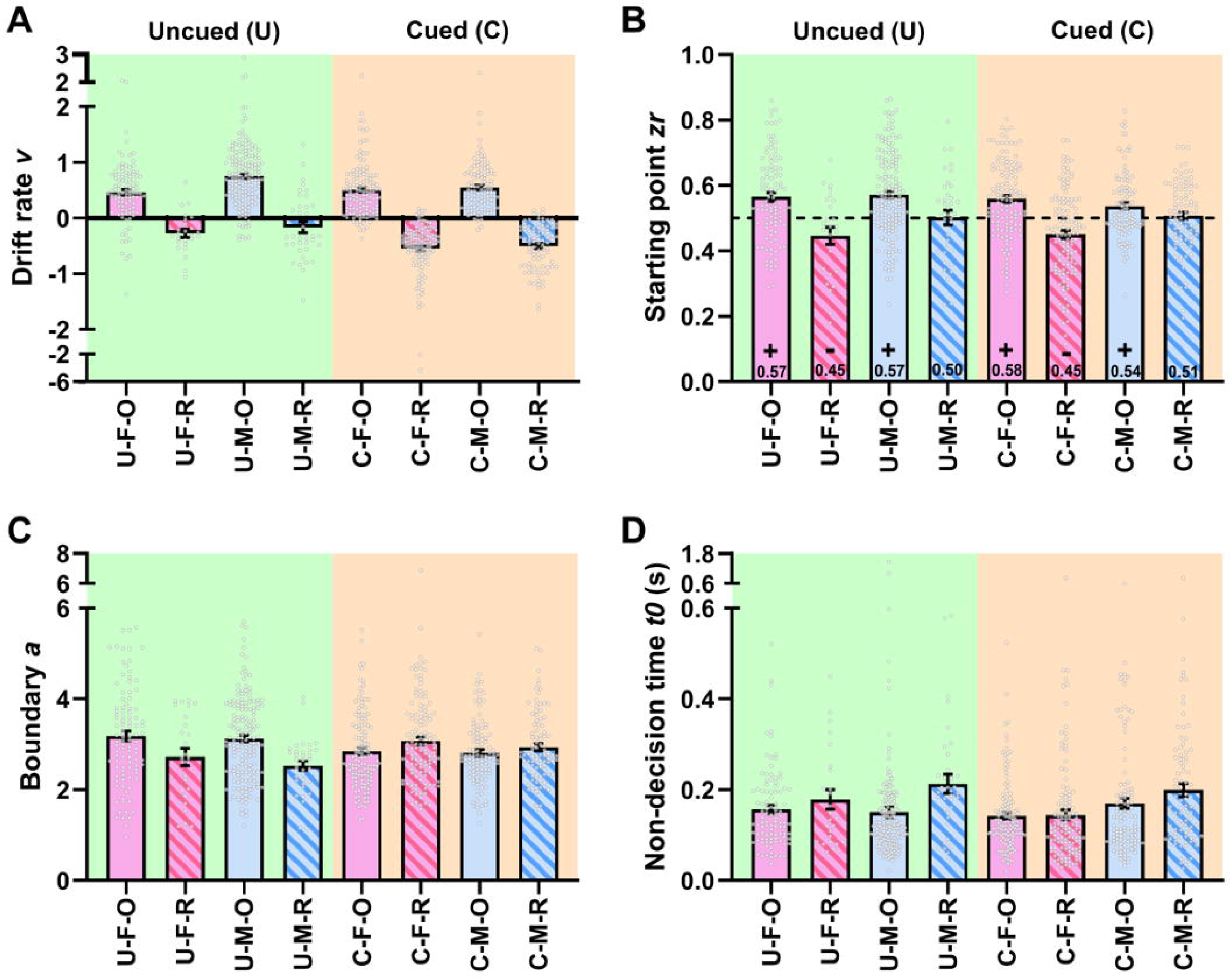
Diffusion model parameters in optimal and risk-preferring rats performing the cued and uncued rat gambling task. Diffusion model parameters derived from rat gambling task (rGT) performance, shown for optimal and risk-preferring rats across cued and uncued task variants and separated by sex. (A) Drift rate: Females exhibited more negative drift rates overall than males. As expected, optimal rats showed positive drift rates toward the optimal decision boundary, whereas risk-preferring rats showed negative drift rates toward the risky boundary. Drift rates were generally more negative on the cued task, driven primarily by risk-preferring animals. (B) Starting point: Optimal rats showed starting-point biases toward optimal choices, whereas risk-preferring rats exhibited biases toward risky choices. This effect was driven by females as male risk-preferring rats displayed neutral starting points regardless of task condition. (C) Boundary separation: The pattern for this measure was dependent on task variant: on the uncued task, optimal rats had wider boundaries, whereas on the cued task risk-preferring rats exhibited wider boundaries than optimal rats. This effect was consistent across sexes. (D) Non-decision time: Risk-preferring rats showed longer non-decision times than optimal rats, and males exhibited longer non-decision times than females. Data are presented as mean ± SEM, with individual data points overlaid. Significant effects are not displayed on this figure for clarity.

#### RP/OPT

As expected, drift rates were positive (towards the optimal boundary) for optimal rats, and negative (towards the risky boundary) for risk-preferring animals (risk preference: F_1,768_ = 546.963, p < 0.001). For risk-preferring rats only, drift rates were more negative on the cued compared to the uncued task (task*risk preference: F_1,768_ = 4.990, p = 0.026). Furthermore, drift rates were more negative overall for the cued task (task: F_1,768_ = 32.015, p < 0.001).

#### PchoiceX

Drift rates were most positive for Pchoice2 rats (P-choice category: F_2,764_ = 247.735, ps < 0.004; Figure 5A). For both Pchoice3 and Pchoice 4 only, drift rates were more negative on the cued task compared to uncued (task*cluster: F_2,764_ = 5.890, p = 0.003, ps < 0.004). Within P2-preferring rats, males had more positive drift rates (towards the optimal boundary) on the uncued task (task*sex: F_1,406_ = 7.988, p = 0.005, uncued vs cued for males: p = 0.002), and these were also more positive compared to females on the uncued task (p < 0.001). This echoes the findings above, showing that animals which preferred the risky options had more negative drift rates on the cued task.

**Figure 5.**
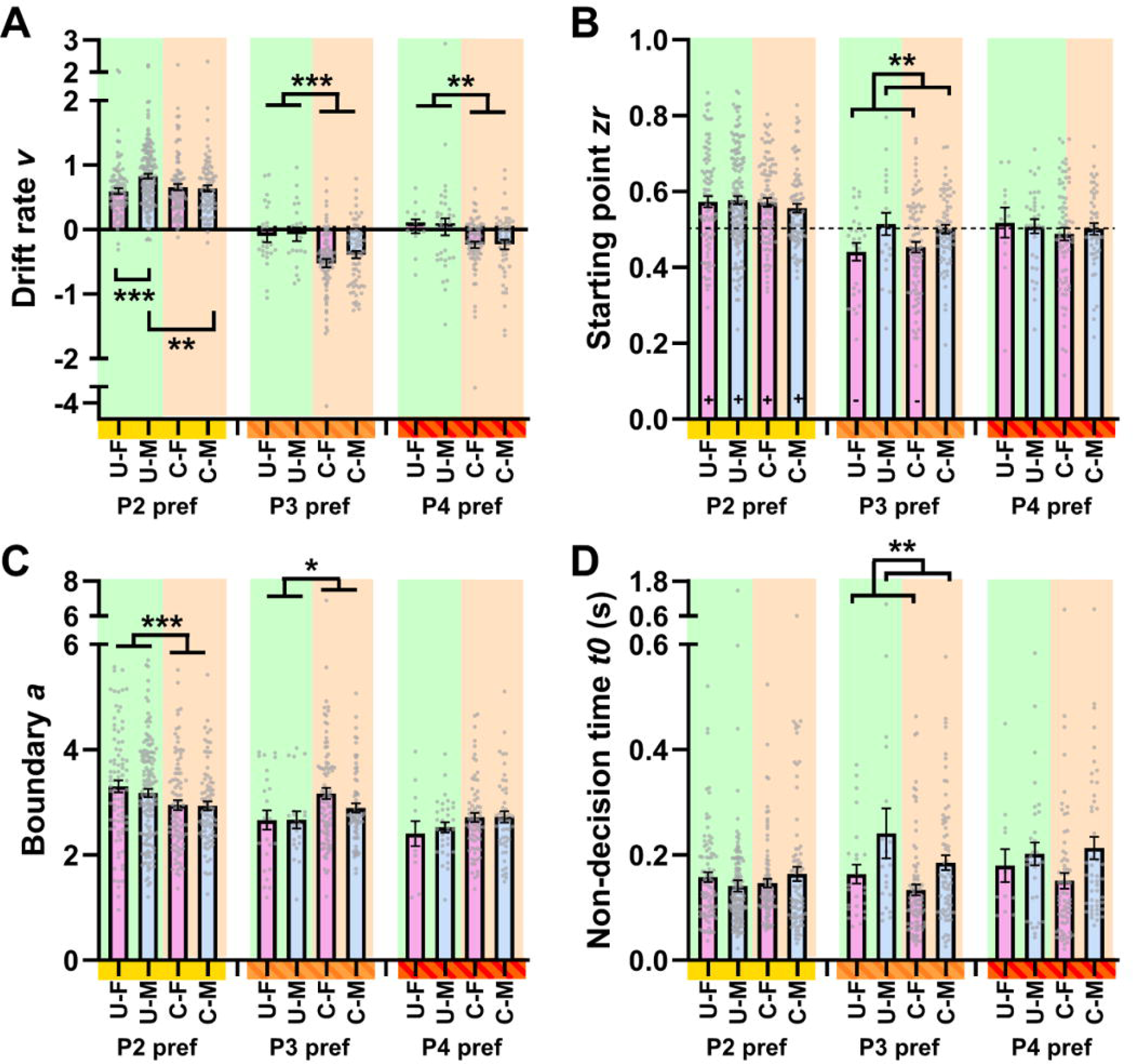
Diffusion model parameters in rats classified according to P-choice category. Diffusion model parameters derived from rat gambling task (rGT) performance, grouped according to Pchoice category (Pchoice2, Pchoice3, or Pchoice4) and separated by task variant and sex. (A) Drift rate: Pchoice2 rats had the most positive drift rates, indicating stronger evidence accumulation toward optimal choices. Drift rates were more negative on the cued task in both Pchoice3 and Pchoice4 rats. Within the Pchoice2 group, males on the uncued task showed positive drift rates compared with females and with males performing the cued task. (B) Starting point: Starting points differed across P-choice categories, with Pchoice2 rats displaying the strongest positive bias toward optimal options. In contrast, female Pchoice3 rats had starting points biased toward risky choices, whereas male Pchoice3 rats showed no bias. Starting points were neutral in Pchoice4 rats regardless of sex or task condition. (C) Boundary separation: Pchoice2 rats had wider decision boundaries than the other groups. Boundary separation was reduced on the cued task in Pchoice2 rats but increased on this task in Pchoice3 rats. (D) Non-decision time: Pchoice4 rats exhibited longer non-decision times than Pchoice2 rats. Within the Pchoice3 subgroup, males had longer non-decision times than females, and non-decision times were longer on the uncued than the cued task. Data are presented as mean ± SEM, with individual data points overlaid. *p<0.05, **p<0.01, ***p<0.001.

#### Starting point

##### RP/OPT

Optimal rats had starting points biased (i.e. different from 0.5, which represents neutral starting point) towards optimal choices, whereas risk-preferring rats had starting points biased towards risky choices (risk preference: F_1,768_ = 60.287, p < 0.001; Figure 4B). However, this latter effect was driven by female rats only (sex*risk preference: F_1,768_ = 9.338, p = 0.002). Male risk-preferring rats did not have biased starting points in either task (one-sample t-tests: ps > 0.518).

##### P-choiceX

Starting points also differed across subgroups (P-choice category: F_2,764_ = 34.968, p < 0.001), with Pchoice2 rats having the most positive, and significantly positive biased starting points (ps < 0.001 for comparisons against Pchoice3 and Pchoice4; one sample t-tests: ps < 0.001, Figure 5B). In Pchoice3 rats only (sex*cluster: F_2,764_ = 3.891, p = 0.021), female rats had negative starting points biased towards risky choices (male vs female: p = 0.003; one sample t-tests: ps < 0.029; Figure 5B), whereas male Pchoice3 rats did not have biased starting points (one sample t-tests: ps > 0.635). All starting points were neutral in Pchoice4 rats (one sample t-tests: ps > 0.468; Figure 5B).

### Boundary

#### RP/OPT

Optimal rats have wider boundaries than risk-preferring rats (risk preference: F_1,768_ = 4.125, p = 0.021; Figure 4C). This is driven by differences on the uncued task only (task*risk preference: F_1,768_ = 21.473, p < 0.001, optimal vs risk-preferring for uncued: p < 0.001), as this pattern is actually reversed on the cued task, with risk-preferring rats having wider decision boundaries than optimal rats (p = 0.035). This pattern is the same across sexes.

#### PchoiceX

Pchoice2 rats had wider decision boundaries than other subgroups (P-choice category: F_2,764_ = 15.135, p < 0.001; ps < 0.004; Figure 5C). Pchoice2 and Pchoice3 rats showed opposite patterns between tasks (task*P-choice category: F_2,764_ = 9.699, p < 0.001), with wider decision boundaries on the uncued vs cued task in Pchoice2 rats (p < 0.001), but wider boundaries for the cued vs uncued task in Pchoice3 rats (p = 0.013).

### Non-decision time

#### RP/OPT

Non-decision times were longer in risk-preferring rats (risk preference: F_1,768_ = 8.140, p = 0.004), and longer for males compared to females (sex: F_1,768_ = 7.323, p = 0.004; Figure 4D).

#### PchoiceX

Non-decision times were longer in Pchoice4 rats, compared to Pchoice2 rats only (P-choice category: F_2,764_ = 5.355, p = 0.005, P4 vs P2: p = 0.012; Figure 5D). Within the Pchoice3 subgroup (sex*cluster: F_2,764_ = 4.362, p = 0.013), males had longer non-decision times than females (p = 0.003). Also, within Pchoice3, non-decision times were longer on the uncued task compared to cued (task*sex: F_1,198_ = 4.580, p = 0.034).

## Discussion

Using an unusually large dataset of over 750 rats, this study provides a comprehensive behavioral and computational analysis of risky decision-making on the rat gambling task (rGT). By comparing traditionally used decision scores with unsupervised clustering to create P-choice–based subgroups, combined with diffusion decision modeling, we demonstrate that win-paired cues not only increase the prevalence of risk-preferring behavior but also fundamentally alter the cognitive processes governing choice across sexes and decision-making phenotypes. These findings help to clarify how cue-driven risk escalation emerges, reveal dissociable mechanisms underlying impulsive versus deliberative risky decisions, and highlight subtle but meaningful sex differences that are often obscured in smaller samples.

### Cue-induced risky decision making is robust, pervasive, and not sex-specific

Consistent with prior reports (e.g. Barrus & Winstanley, 2016), the addition of win-paired cues markedly increased the proportion of rats classified as risk-preferring. Importantly, the magnitude of this effect was equivalent in males and females, demonstrating that cues exert a powerful influence on decision-making regardless of biological sex. Beyond replicating this finding, the present results extend it in two critical ways.

First, riskier choice was evident within both optimal and risk-preferring rats. Decision scores became more negative on the cued task even in rats that remained classified as optimal, as well as in rats already inclined toward risky choice. A similar pattern emerged when rats were stratified by P-choice preference: P2-, P3-, and P4-preferring rats all exhibited more negative decision scores on the cued rGT. Therefore, cues do not simply recruit a subset of animals into a risk-preferring category; rather, they shift the entire population towards riskier choices.

Second, diffusion modeling revealed that this behavioral shift is accompanied by changes in latent cognitive parameters. Drift rates were more negative on the cued task overall, indicating that evidence accumulation was biased toward risky options when rewards were signaled by salient cues. This was especially pronounced in risk-preferring rats and in P3-and P4-preferring subgroups, reinforcing the conclusion that cue-induced risky choice reflects altered option evaluation dynamics rather than just changes in response vigor or motivation.

### Sex differences emerge in how cues alter risky choice

Although males and females showed comparable proportions of risk-preferring behavior at the aggregate level, several more in-depth analyses revealed sex-specific differences in how reward cues influenced choice patterns and underlying decision processes. Most strikingly, females showed a pronounced reduction in P1 choice on the cued task, accompanied by increased selection of the riskiest option (P4), regardless of whether they were classified as optimal or risk-preferring. In contrast, cue-driven changes in males were characterized more by increased P3 selection, with divergent effects on P4 depending on baseline risk preference.

These findings suggest that cues may differentially alter learning or valuation strategies across sexes. One possibility is that females are more sensitive to the relative salience or perceived value of cue-paired outcomes, leading to a disproportionate devaluation of low-reward options such as P1. This interpretation aligns with recent reinforcement learning computational modelling, suggesting sex differences in how reward-predictive cues influence value learning and belief updating (Hales et al., 2025). Notably, in the absence of cues, P-choice patterns in optimal males and females were nearly identical, indicating that these sex differences are not intrinsic to baseline task performance but emerge specifically in response to cue presence.

Diffusion model parameters further support the idea of sex-dependent cognitive mechanisms. Drift rates were more negative overall in females, and female P2-preferring rats exhibited particularly negative drift rates on the uncued task compared to males. Moreover, only female risk-preferring rats showed starting point biases toward risky choices, whereas male risk-preferring rats exhibited neutral starting points. This dissociation suggests that risky choice in females may be driven partly by pre-decisional biases or expectations, whereas in males it may rely more heavily on trial-by-trial evidence accumulation or other processes. Such distinctions would be difficult to infer from choice behavior alone, underscoring the value of computational modeling, and may imply divergent neural mechanisms mediate risky choice across the sexes. In support of this conclusion, recent work suggests that chemogenetic manipulation of dopaminergic projections arising from the VTA and cholinergic interneurons within the nucleus accumbens have opposing effects on cue-induced risky choice in male and female rats (Hynes et al., 2020; 2021; 2024; 2025).

### Risky decisions are not uniformly impulsive and often involve deliberation

A central contribution of this study is the demonstration that risky decision-making on the rGT cannot be directly equated with impulsivity. While risk-preferring rats made more premature responses than optimal decision-makers, risky choices were generally associated with longer, not shorter, response latencies. Risk-preferring rats were slower than optimal rats when selecting risky options, and P3-and P4-preferring animals were slower than P2-preferring rats to choose disadvantageous options. These findings argue against a simple speed–accuracy trade-off in which risky choices arise rashly, through inadequate deliberation.

Diffusion modeling offers a mechanistic explanation for this apparent paradox. Decision boundary separation showed a U-shaped relationship with decision score: both highly optimal and highly risk-preferring rats exhibited wider boundaries, indicating greater response caution. Notably, this pattern depended on task context. Optimal rats had wider boundaries on the uncued task, whereas risk-preferring rats had wider boundaries on the cued task. A similar dissociation was observed in the P-choice analysis, where P2-preferring rats had the widest boundaries on the uncued rGT, while P3-preferring rats exhibited wider boundaries when rewards were cued. This is consistent with a growing body of work showing that suboptimal or risky decisions are not necessarily impulsive, but can instead arise from biased evidence accumulation processes. In value-based decision-making tasks, individuals may accumulate evidence more slowly yet systematically toward disadvantageous options when subjective value representations are distorted (Shahar et al., 2019; Pedersen et al., 2021). Similarly, altered drift rates have been linked to maladaptive reward processing in clinical populations, even in the presence of intact or increased response caution (Moustafa et al., 2015; Pitliya et al., 2022). These findings support the idea that the deliberative risky choices observed here, particularly on the cued task, may reflect changes in value integration rather than a simple failure of inhibitory control.

Cues shifted decision-making towards risky choices primarily by altering evidence accumulation, rather than uniformly altering decision caution. From a mechanistic perspective, these findings are consistent with theoretical and neurophysiological accounts of the DDM, in which drift rate reflects the accumulation of value-based evidence and boundary separation represents strategic control over decision thresholds. Neural correlates of these processes have been identified in distributed cortical and subcortical circuits, suggesting that distinct systems likely modulate evidence accumulation and response caution (Mueller et al., 2017; Imani et al., 2023).

Together, these results suggest that risky choices can arise through at least two distinct cognitive routes. In one route, more prevalent on the uncued task, risky choices may be driven by impulsive responding characterized by narrow boundaries, negative drift rates and high premature response rates. In another route, particularly evident on the cued task, risky choices appear more deliberative, with wider boundaries and slower response times, despite an overall bias toward risky outcomes. A similar pattern has been reported in humans, whereby participants took longer to choose between two gambles when the outcomes were accompanied by audiovisual cues (Baumann et al., 2025). This divergence in the cognitive mechanisms underlying risky choice may help explain why pharmacological or experimental manipulations selectively improve decision-making on one task variant, or have opposing effects on the cued versus uncued rGT. For example, the noradrenaline reuptake inhibitor atomoxetine, the cholinergic muscarinic antagononist scopolamine, and the serotonin 2C receptor antagonist SB 242084 all decrease risky choice, but only when rewards are cued (Silveira et al., 2015; 2016; Adams et al., 2017; Betts et al., 2021; Chernoff et al., 2021). The addition of win-paired cues to the rGT therefore seems to fundamentally shift the neurocognitive circuitry involved in the decision-making process.

### Dissociating motor impulsivity from risky choice

The present findings also clarify the relationship between motor impulsivity and risky choice. Premature responding correlated negatively with drift rate, boundary separation and non-decision time, consistent with the interpretation that impulsive actions generally track faster, less constrained decision processes. Impulsive adolescents with borderline personality disorder also exhibited lower decision-boundaries in an emotional interference task (Forester et al., 2026), while stop-signal reaction times were more strongly correlated with non decision time in humans performing the stop signal task (White et al., 2014). However, premature responding was not uniformly elevated by win-paired cues. Instead, cue-induced increases in premature responding was driven entirely by optimal or P2-preferring rats - animals that are typically less impulsive.

This dissociation aligns with computational accounts demonstrating that different facets of impulsivity map onto distinct DDM parameters. Reduced boundary separation has been associated with impulsive action and diminished response caution, whereas drift rate biases are more closely linked to value-based impulsivity and altered reward sensitivity (Moustafa et al., 2015; Johnson et al., 2017; Sharma & Khan, 2018; Pitliya et al., 2022). The present findings extend this framework by showing that motor impulsivity and risky decision-making can diverge not only behaviorally but also at the level of underlying computational mechanisms.

Risk-preferring rats, although more impulsive overall, made comparable numbers of premature responses regardless of cue presence, echoing findings from the choice rGT paradigm (Hales et al. 2025), and suggesting that the elevated impulsivity associated with risk preference is a stable trait rather than a cue-induced state. Thus, win-paired cues appear to drive impulsive responding primarily in rats that would otherwise exert greater motor control, and such cue-driven impulse control failures may depend on distinct cognitive processes or neurobiology. In support of this hypothesis, pharmacological modulation of premature responding can diverge across the cued and uncued task. For example, the dopamine D2/3 agonist ropinirole caused a more robust and long-lasting increase in premature responding on the cued rGT (Tremblay et al. 2019), while the nicotinic receptor antagonist mecamylamine decreased premature responding on the cued rGT only (Silveira et al., 2015; Betts et al., 2021).

### Task engagement, motivation and sex differences

Advantageous decision-making was associated with greater task engagement, as reflected by more trials completed and fewer time-out penalties. P2-preferring rats completed the most trials, and optimal decision-makers outperformed risk-preferring rats in this regard. Sex differences were also evident: males completed more trials and made fewer omissions overall, particularly on the uncued task. While these effects could reflect differences in motivation or physical factors such as body size, they may also relate to sex-dependent strategies for balancing reward pursuit and effort expenditure. Importantly, these engagement differences did not straightforwardly map onto risk preference, reinforcing the notion that risky choice is not just a result of reduced task involvement.

### Implications for classification strategies and translational relevance

By comparing decision score–based classification with P-choice clustering, this study demonstrates that both approaches capture meaningful but partially distinct aspects of rGT behavior. The convergence of findings across classification schemes strengthens confidence in the core conclusions. At the same time, diffusion modeling revealed mechanistic heterogeneity even within apparently similar behavioral groups, highlighting the limitations of relying on any single metric to characterize complex decision-making.

From a translational perspective, these results underscore the importance of distinguishing between impulsive and deliberative forms of maladaptive choice, particularly in contexts involving salient cues, such as substance use disorders or gambling disorder. The finding that cues can promote risk-taking through deliberative processes challenges simplistic models in which cue-reactivity negatively effects cognitive control, and instead suggests that cues may alter reward valuation in a more nuanced manner. Sex-specific mechanisms further emphasize the need for inclusive, large-scale approaches when modeling psychiatric vulnerability.

### Conclusions

In summary, this large-scale behavioral and computational analysis demonstrates that reward-paired cues robustly and pervasively bias decision-making toward risk on the rGT, across sexes and decision-making phenotypes. Crucially, risky choice emerges from multiple cognitive pathways that differ in their reliance on impulsivity, deliberation and pre-decisional bias. By integrating diffusion modeling with fine-grained behavioral analyses, the present study advances our understanding of how cues, sex and individual traits interact to shape risky decision-making, and provides a framework for interpreting mixed outcomes in both basic and translational research.

## Acknowledgements

This work took place at a UBC campus situated on the traditional, ancestral, and unceded land of the x□məθk□əy□əm (Musqueam) People. We acknowledge and are grateful for their stewardship of this land for thousands of years.

## Author contributions

CAH and CAW developed the hypotheses to be tested, discussed how to interpret the results, and co-wrote the manuscript. CAH ran all the modeling and statistical analyses.

## References

1. Adams, W. K., Barkus, C., Ferland, J. N., Sharp, T., & Winstanley, C. A. (2017). Pharmacological evidence that 5-HT_2C_ receptor blockade selectively improves decision making when rewards are paired with audiovisual cues in a rat gambling task. Psychopharmacology, 234(20), 3091–3104. 10.1007/s00213-017-4696-4

2. Adams, W. K., Vonder Haar, C., Tremblay, M., Cocker, P. J., Silveira, M. M., Kaur, S., Baunez, C., & Winstanley, C. A. (2017). Deep-Brain Stimulation of the Subthalamic Nucleus Selectively Decreases Risky Choice in Risk-Preferring Rats. eNeuro, 4(4), ENEURO.0094-17.2017. 10.1523/ENEURO.0094-17.2017

3. Barrus, M. M., & Winstanley, C. A. (2016). Dopamine D3 Receptors Modulate the Ability of Win-Paired Cues to Increase Risky Choice in a Rat Gambling Task. The Journal of neuroscience : the official journal of the Society for Neuroscience, 36(3), 785–794. 10.1523/JNEUROSCI.2225-15.2016

4. Barrus, M. M., & Winstanley, C. A. (2017). Cued Rat Gambling Task. Bio-protocol, 7(3), e2118. 10.21769/BioProtoc.2118

5. Baumann, G., Fischer, R., Reinhard, I., Hoffmann, S., Kiefer, F., Leménager, T., & Bach, P. (2025). Investigating Decision-Making under Risk in Pathological Gambling Using a Virtual Slot Machine: A Pilot Eye-Tracking Study. European addiction research, 31(5), 308–324. 10.1159/000547742

6. Bechara, A., Damasio, A. R., Damasio, H., & Anderson, S. W. (1994). Insensitivity to future consequences following damage to human prefrontal cortex. Cognition, 50(1-3), 7–15. 10.1016/0010-0277(94)90018-3

7. Betts, G. D., Hynes, T. J., & Winstanley, C. A. (2021). Pharmacological evidence of a cholinergic contribution to elevated impulsivity and risky decision-making caused by adding win-paired cues to a rat gambling task. Journal of psychopharmacology (Oxford, England), 35(6), 701–712. 10.1177/0269881120972421

8. Brevers, D., Bechara, A., Cleeremans, A., & Noël, X. (2013). Iowa Gambling Task (IGT): twenty years after - gambling disorder and IGT. Frontiers in psychology, 4, 665. 10.3389/fpsyg.2013.00665

9. Byrnes, J. P., Miller, D. C., & Schafer, W. D. (1999). Gender differences in risk taking: A meta-analysis. Psychological Bulletin, 125(3), 367–383. 10.1037/0033-2909.125.3.367

10. Chernoff, C. S., Hynes, T. J., & Winstanley, C. A. (2021). Noradrenergic contributions to cue-driven risk-taking and impulsivity. Psychopharmacology, 238(7), 1765–1779. 10.1007/s00213-021-05806-x

11. Cross, C. P., Copping, L. T., & Campbell, A. (2011). Sex differences in impulsivity: a meta-analysis. Psychological bulletin, 137(1), 97–130. 10.1037/a0021591

12. Ferland, J. N., & Winstanley, C. A. (2017). Risk-preferring rats make worse decisions and show increased incubation of craving after cocaine self-administration. Addiction biology, 22(4), 991– 1001. 10.1111/adb.12388

13. Forester, G., Richson, B. N., Reilly, E. E., Anderson, L. M., Wonderlich, S. A., & Schaefer, L. M. (2026). Longitudinal analysis of decision-making deficits in binge-eating disorders using drift diffusion modeling. Appetite, 222, 108497. Advance online publication. 10.1016/j.appet.2026.108497

14. Hales, C. A., Houghton, C. J., & Robinson, E. S. J. (2017). Behavioural and computational methods reveal differential effects for how delayed and rapid onset antidepressants effect decision making in rats. European neuropsychopharmacology : the journal of the European College of Neuropsychopharmacology, 27(12), 1268–1280. 10.1016/j.euroneuro.2017.09.008

15. Hales, C. A., Robinson, E. S., & Houghton, C. J. (2016). Diffusion Modelling Reveals the Decision Making Processes Underlying Negative Judgement Bias in Rats. PloS one, 11(3), e0152592. 10.1371/journal.pone.0152592

16. Hales, C. A., Silveira, M. M., Calderhead, L., Mortazavi, L., Hathaway, B. A., & Winstanley, C. A. (2024). Insight into differing decision-making strategies that underlie cognitively effort-based decision making using computational modeling in rats. Psychopharmacology, 241(5), 947–962. 10.1007/s00213-023-06521-5

17. Hales, C. A., Hrelja, K. M., Ansary, S., Chong, E., Russell, B., & Winstanley, C. A. (2025). Most rats prefer gambling opportunities featuring win-paired cues that drive risky choice: Synergistic interactions between choice of and choice during the cued rat gambling task. Brain and neuroscience advances, 9, 23982128251352235. 10.1177/23982128251352235

18. Hathaway, B. A., Schumacher, J. D., Hrelja, K. M., & Winstanley, C. A. (2021). Serotonin 2C Antagonism in the Lateral Orbitofrontal Cortex Ameliorates Cue-Enhanced Risk Preference and Restores Sensitivity to Reinforcer Devaluation in Male Rats. eNeuro, 8(6), ENEURO.0341-21.2021. 10.1523/ENEURO.0341-21.2021

19. Hultman, C., Tjernström, N., Vadlin, S., Rehn, M., Nilsson, K. W., Roman, E., & Åslund, C. (2022). Exploring decision-making strategies in the Iowa gambling task and rat gambling task. Frontiers in behavioral neuroscience, 16, 964348. 10.3389/fnbeh.2022.964348

20. Hynes, T. J., Chernoff, C. S., Hrelja, K. M., Tse, M. T. L., Avramidis, D. K., Lysenko-Martin, M. R., Calderhead, L., Kaur, S., Floresco, S. B., & Winstanley, C. A. (2024). Win-Paired Cues Modulate the Effect of Dopamine Neuron Sensitization on Decision Making and Cocaine Self-administration: Divergent Effects Across Sex. Biological psychiatry, 95(3), 220–230. 10.1016/j.biopsych.2023.08.021

21. Hynes, T. J., Chernoff, C. S., Hrelja, K., Li, A., Betts, G. D., Calderhead, L. S., & Winstanley, C. A. (2025). Ventral Striatal Cholinergic Interneurons Regulate Decision-Making or Motor Impulsivity Differentially across Learning and Biological Sex. The Journal of neuroscience : the official journal of the Society for Neuroscience, 45(49), e0764252025. 10.1523/JNEUROSCI.0764-25.2025

22. Hynes, T. J., Hrelja, K. M., Hathaway, B. A., Hounjet, C. D., Chernoff, C. S., Ebsary, S. A., Betts, G. D., Russell, B., Ma, L., Kaur, S., & Winstanley, C. A. (2021). Dopamine neurons gate the intersection of cocaine use, decision making, and impulsivity. Addiction biology, 26(6), e13022. 10.1111/adb.13022

23. Imani, E., Radkani, S., Hashemi, A., Harati, A., Pourreza, H., & Moazami Goudarzi, M. (2023). Distributed Coding of Evidence Accumulation across the Mouse Brain Using Microcircuits with a Diversity of Timescales. eNeuro, 10(11), ENEURO.0282-23.2023. 10.1523/ENEURO.0282-23.2023

24. Johnson, D. J., Hopwood, C. J., Cesario, J., & Pleskac, T. J. (2017). Advancing research on cognitive processes in social and personality psychology: A hierarchical drift diffusion model primer. Social Psychological and Personality Science,8 (4), 413–423.

25. Langdon, A. J., Hathaway, B. A., Zorowitz, S., Harris, C. B. W., & Winstanley, C. A. (2019). Relative insensitivity to time-out punishments induced by win-paired cues in a rat gambling task. Psychopharmacology, 236(8), 2543–2556. 10.1007/s00213-019-05308-x

26. Lindberg, F. A., Kagios, C., Tjernström, N., & Roman, E. (2025). Individual differences in training time in the rat gambling task are unrelated to subsequent decision-making strategies. Frontiers in psychiatry, 16, 1490196. 10.3389/fpsyt.2025.1490196

27. Moustafa, A. A., Kéri, S., Somlai, Z., Balsdon, T., Frydecka, D., Misiak, B., & White, C. (2015). Drift diffusion model of reward and punishment learning in schizophrenia: Modeling and experimental data. Behavioural brain research, 291, 147–154. 10.1016/j.bbr.2015.05.024

28. Mueller, C. J., White, C. N., & Kuchinke, L. (2017). Electrophysiological correlates of the drift diffusion model in visual word recognition. Human brain mapping, 38(11), 5616–5627. 10.1002/hbm.23753

29. Nelder, J. A., & Mead, R. (1965) A simplex method for function minimization. The Computer Journal 7(4): 308–313. 10.1093/comjnl/7.4.308

30. Orsini, C. A., & Setlow, B. (2017). Sex differences in animal models of decision making. Journal of neuroscience research, 95(1-2), 260–269. 10.1002/jnr.23810

31. Orsini, C. A., Willis, M. L., Gilbert, R. J., Bizon, J. L., & Setlow, B. (2016). Sex differences in a rat model of risky decision making. Behavioral neuroscience, 130(1), 50–61. 10.1037/bne0000111

32. Pedersen, M. L., Ironside, M., Amemori, K. I., McGrath, C. L., Kang, M. S., Graybiel, A. M., Pizzagalli, D. A., & Frank, M. J. (2021). Computational phenotyping of brain-behavior dynamics underlying approach-avoidance conflict in major depressive disorder. PLoS computational biology, 17(5), e1008955.

33. Pitliya, R. J., Nelson, B. D., Hajcak, G., & Jin, J. (2022). Drift-Diffusion Model Reveals Impaired Reward-Based Perceptual Decision-Making Processes Associated with Depression in Late Childhood and Early Adolescent Girls. Research on child and adolescent psychopathology, 50(11), 1515–1528. 10.1007/s10802-022-00936-y

34. Rivalan, M., Valton, V., Seriès, P., Marchand, A. R., & Dellu-Hagedorn, F. (2013). Elucidating poor decision-making in a rat gambling task. PloS one, 8(12), e82052. 10.1371/journal.pone.0082052

35. Shahar, N., Hauser, T. U., Moutoussis, M., Moran, R., Keramati, M., NSPN consortium, & Dolan, R. J. (2019). Improving the reliability of model-based decision-making estimates in the two-stage decision task with reaction-times and drift-diffusion modeling. PLoS computational biology, 15(2), e1006803. 10.1371/journal.pcbi.1006803

36. Sharma, S. N., & Khan, A. (2018). Interval timing predicts impulsivity in intertemporal choice: combined behavioral and drift-diffusion model evidence. Journal of Cognitive Psychology, 30(8), 816–831. 10.1080/20445911.2018.1539002

37. Shen, L., Hu, Yx., Lv, Qy. et al. Using hierarchical drift diffusion models to elucidate computational mechanisms of reduced reward sensitivity in adolescent major depressive disorder. BMC Psychiatry 24, 933 (2024). 10.1186/s12888-024-06353-3

38. Silveira, M. M., Malcolm, E., Shoaib, M., & Winstanley, C. A. (2015). Scopolamine and amphetamine produce similar decision-making deficits on a rat gambling task via independent pathways. Behavioural brain research, 281, 86–95. 10.1016/j.bbr.2014.12.029

39. Silveira, M. M., Murch, W. S., Clark, L., & Winstanley, C. A. (2016). Chronic atomoxetine treatment during adolescence does not influence decision-making on a rodent gambling task, but does modulate amphetamine’s effect on impulsive action in adulthood. Behavioural pharmacology, 27(4), 350–363. 10.1097/FBP.0000000000000203

40. Tjernström, N., & Roman, E. (2022). Individual strategies in the rat gambling task are related to voluntary alcohol intake, but not sexual behavior, and can be modulated by naltrexone. Frontiers in psychiatry, 13, 931241. 10.3389/fpsyt.2022.931241

41. Tremblay, M., Barrus, M. M., Cocker, P. J., Baunez, C., & Winstanley, C. A. (2019). Increased motor impulsivity in a rat gambling task during chronic ropinirole treatment: potentiation by win-paired audiovisual cues. Psychopharmacology, 236(6), 1901–1915. 10.1007/s00213-019-5173-z

42. van den Bos, R., Homberg, J., & de Visser, L. (2013). A critical review of sex differences in decision-making tasks: focus on the Iowa Gambling Task. Behavioural brain research, 238, 95–108. 10.1016/j.bbr.2012.10.002

43. Vonder Haar, C., Frankot, M. A., Reck, A. M., Milleson, V., & Martens, K. M. (2022). Large-N Rat Data Enables Phenotyping of Risky Decision-Making: A Retrospective Analysis of Brain Injury on the Rodent Gambling Task. Frontiers in behavioral neuroscience, 16, 837654. 10.3389/fnbeh.2022.837654

44. Voss, A., & Voss, J. (2007). Fast-dm: a free program for efficient diffusion model analysis. Behavior research methods, 39(4), 767–775. 10.3758/bf03192967

45. Voss, A., & Voss, J. (2008) A fast numerical algorithm for the estimation of diffusion model parameters. Journal of Mathematical Psychology 52(1): 1–9. 10.1016/j.jmp.2007.09.005

46. Voss, A., Voss, J., & Klauer, K. C. (2010). Separating response-execution bias from decision bias: arguments for an additional parameter in Ratcliff’s diffusion model. The British journal of mathematical and statistical psychology, 63(Pt 3), 539–555. 10.1348/000711009X477581

47. Voss A, Voss J, Lerche V (2015) Assessing cognitive processes with diffusion model analyses: A tutorial based on fast-dm-30. Frontiers in Psychology 6: 336. 10.3389/fpsyg.2015.00336

48. White, C. N., Congdon, E., Mumford, J. A., Karlsgodt, K. H., Sabb, F. W., Freimer, N. B., London, E. D., Cannon, T. D., Bilder, R. M., & Poldrack, R. A. (2014). Decomposing decision components in the stop-signal task: a model-based approach to individual differences in inhibitory control. Journal of cognitive neuroscience, 26(8), 1601–1614. 10.1162/jocn_a_00567

49. Zeeb, F. D., Robbins, T. W., & Winstanley, C. A. (2009). Serotonergic and dopaminergic modulation of gambling behavior as assessed using a novel rat gambling task. Neuropsychopharmacology : official publication of the American College of Neuropsychopharmacology, 34(10), 2329–2343. 10.1038/npp.2009.62

