## Supplementary Material for "Applying drift diffusion models to rat gambling task data reveals divergent cognitive mechanisms underlying risky choice"

**Author names:** Claire A. Hales, Catharine A. Winstanley

**Affiliations:** *Department of Psychology, Djavad Mowafaghian Centre for Brain Health, University of British Columbia, Vancouver, BC, Canada*

**Corresponding author:** Correspondence should be addressed to Dr. Claire A. Hales or Dr. Catharine A. Winstanley

*Correspondence address:* Djavad Mowafaghian Centre for Brain Health, Department of Psychology, 2215 Wesbrook Mall, Vancouver, BC, V6T 1Z3, Canada

|  | **Cluster number** | | | | |
| --- | --- | --- | --- | --- | --- |
| **Optimal cluster criterion** | **2** | **3** | **4** | **5** | **6** |
| Calinski-Harabasz | 2057 | 2556 | 2883 | **2966** | 2784 |
| Davies-Bouldin | **0.5466** | 0.5717 | 0.6220 | 0.6837 | 0.7646 |
| Gap | 0.2253 | 0.4840 | 0.7121 | **0.8425** | 0.7932 |
| Silhouette | **0.7805** | 0.7291 | 0.7047 | 0.6777 | 0.6342 |

**Table 1** – *Optimal cluster criterion value comparison.*

Four methods to determine optimal cluster number (using evalclusters function in Matlab) were compared for k-means clustering run on stable acquisition rGT data. Bold font denotes the optimal cluster number found for each method. Calinkski-Harabasz criterion: this method is also known as the variance ratio criterion. Well-defined clusters have large between-cluster variance and small within-cluster variance; therefore, the optimal number of clusters corresponds to the highest Calinski-Harabasz value. Davies-Bouldin criterion: this method is based on a ratio of within-cluster and between-cluster distances, and so the optimal solution has the smallest Davies-Bouldin value. Gap value criterion: values correspond to: *ExpectedLogW – LogW*, where *W* is the within-cluster dispersion, *ExpectedLogW* is found through Monte Carlo sampling of a reference distribution, and *LogW* is calculated from the input data. The solution with the largest gap value (within a tolerance range) corresponds to the optimal solution. Silhouette criterion: The silhouette value for each datapoint is a measure similarity between that point and other points in the same cluster, compared to points in other clusters. Optimal clustering corresponds to the solution where most datapoints have large silhouette values.

| Measure | Task | Sex | Cluster | | | | |
| --- | --- | --- | --- | --- | --- | --- | --- |
|  |  |  | **Highly optimal** | **Mid optimal** | **Low optimal** | **Low risky** | **Highly risky** |
| Total trials | **Uncued** | **F** | 110.65±3.95 | 93.00±4.76 | 82.54±3.56 | 68.46±4.38 | 59.14±4.78 |
|  |  | **M** | 137.93±2.39 | 108.15±2.86 | 92.91±3.21 | 79.21±2.38 | 70.50±3.29 |
|  | **Cued** | **F** | 116.74±4.77 | 99.86±3.56 | 90.84±2.73 | 78.56±2.05 | 74.12±2.11 |
|  |  | **M** | 131.25±3.33 | 110.06±4.61 | 97.58±4.45 | 78.52±2.50 | 74.17±1.81 |
| Risk score | **Uncued** | **F** | 87.01±1.24 | 51.21±1.88 | 13.10±2.18 | -23.18±2.65 | -63.25±3.68 |
|  |  | **M** | 88.32±0.89 | 50.11±1.60 | 13.45±1.76 | -23.36±2.76 | -76.26±3.73 |
|  | **Cued** | **F** | 89.40±1.20 | 50.20±1.33 | 13.89±1.24 | -24.90±1.80 | -75.89±1.80 |
|  |  | **M** | 85.41±1.38 | 46.26±1.81 | 14.44±1.90 | -24.38±1.91 | -72.78±2.05 |
| % P1 | **Uncued** | **F** | 8.44±1.09 | 15.70±2.31 | 12.47±2.95 | 18.00±3.37 | 6.73±2.19 |
|  |  | **M** | 12.11±1.30 | 14.88±1.78 | 9.90±2.02 | 14.10±2.82 | 5.36±1.33 |
|  | **Cued** | **F** | 4.04±0.78 | 10.09±2.13 | 7.45±1.31 | 5.78±1.07 | 4.09±0.56 |
|  |  | **M** | 8.96±1.76 | 12.37±2.57 | 11.75±2.03 | 10.18±1.76 | 4.88±0.78 |
| % P2 | **Uncued** | **F** | 85.22±1.45 | 59.81±2.20 | 44.46±2.87 | 20.78±3.13 | 10.98±1.59 |
|  |  | **M** | 82.42±1.50 | 60.48±1.84 | 46.96±1.99 | 24.28±2.93 | 5.99±1.74 |
|  | **Cued** | **F** | 79.96±3.78 | 59.02±3.33 | 50.46±1.46 | 32.70±1.68 | 8.62±1.38 |
|  |  | **M** | 84.04±2.11 | 61.01±2.84 | 45.74±2.20 | 27.67±1.98 | 8.33±0.91 |
| % P3 | **Uncued** | **F** | 3.72±0.45 | 11.86±1.57 | 19.99±4.10 | 40.71±6.41 | 71.36±5.71 |
|  |  | **M** | 3.21±0.36 | 9.92±1.33 | 19.05±3.55 | 30.11±4.56 | 32.42±9.99 |
|  | **Cued** | **F** | 10.06±2.70 | 14.80±2.47 | 18.94±2.44 | 29.72±3.70 | 61.27±3.90 |
|  |  | **M** | 4.46±0.60 | 13.15±1.87 | 21.64±2.76 | 37.62±3.94 | 59.04±4.27 |
| % P4 | **Uncued** | **F** | 2.63±0.36 | 12.63±1.85 | 23.08±4.04 | 20.51±6.38 | 10.93±5.83 |
|  |  | **M** | 2.26±0.30 | 14.71±1.43 | 24.10±3.53 | 31.18±4.86 | 56.22±10.79 |
|  | **Cued** | **F** | 8.94±1.91 | 16.09±2.01 | 23.15±2.59 | 31.80±3.49 | 26.02±3.63 |
|  |  | **M** | 2.54±0.54 | 13.47±1.99 | 20.87±2.38 | 24.53±3.67 | 27.75±4.35 |
| Optimal choice latency | **Uncued** | **F** | 1.84±0.10 | 1.60±0.12 | 1.65±0.17 | 1.46±0.27 | 1.41±0.22 |
|  |  | **M** | 1.43±0.07 | 1.53±0.10 | 1.58±0.16 | 1.60±0.12 | 1.72±0.22 |
|  | **Cued** | **F** | 1.42±0.08 | 1.49±0.10 | 1.45±0.06 | 1.59±0.09 | 1.42±0.09 |
|  |  | **M** | 1.38±0.10 | 1.62±0.13 | 1.71±0.10 | 2.10±0.15 | 1.47±0.09 |
| Risky choice latency | **Uncued** | **F** | 1.52±0.13 | 1.73±0.16 | 1.55±0.23 | 1.73±0.25 | 1.50±0.26 |
|  |  | **M** | 1.20±0.09 | 1.31±0.09 | 1.67±0.26 | 1.60±0.12 | 1.36±0.12 |
|  | **Cued** | **F** | 1.36±0.12 | 1.52±0.10 | 1.55±0.10 | 1.75±0.11 | 1.72±0.09 |
|  |  | **M** | 1.20±0.13 | 1.59±0.15 | 1.52±0.13 | 2.06±0.15 | 1.60±0.09 |
| % Premature responses | **Uncued** | **F** | 10.70±0.80 | 13.10±1.06 | 16.17±1.86 | 18.72±3.23 | 20.54±2.24 |
|  |  | **M** | 14.25±0.91 | 18.80±1.64 | 22.34±2.66 | 22.63±2.48 | 30.82±3.96 |
|  | **Cued** | **F** | 15.84±1.03 | 16.91±1.51 | 21.34±1.63 | 16.18±1.12 | 20.73±1.12 |
|  |  | **M** | 16.37±1.48 | 19.98±1.72 | 24.06±2.13 | 20.25±2.00 | 25.03±1.68 |
| % Omissions | **Uncued** | **F** | 3.06±0.46 | 2.89±0.50 | 1.70±0.41 | 4.32±2.14 | 1.90±0.54 |
|  |  | **M** | 1.30±0.17 | 2.02±0.60 | 2.25±0.85 | 1.58±0.55 | 1.28±0.65 |
|  | **Cued** | **F** | 1.63±0.34 | 1.81±0.31 | 2.84±0.45 | 2.49±0.38 | 2.85±0.52 |
|  |  | **M** | 1.07±0.17 | 1.84±0.42 | 1.77±0.36 | 2.68±0.68 | 1.30±0.21 |

**Supplementary Table 2** – *Behavioural measures analysed by cluster*

**Supplementary Table 3** – *Diffusion model parameters by cluster*

| Measure | Task | Sex | Cluster | | | | |
| --- | --- | --- | --- | --- | --- | --- | --- |
|  |  |  | **Highly optimal** | **Mid optimal** | **Low optimal** | **Low risky** | **Highly risky** |
| Drift rate | **Uncued** | **F** | 0.75±0.06 | 0.26±0.06 | 0.19±0.09 | -0.24±0.09 | -0.53±0.13 |
|  |  | **M** | 0.98±0.04 | 0.47±0.05 | 0.23±0.14 | 0.06±0.12 | -0.57±0.14 |
|  | **Cued** | **F** | 0.96±0.06 | 0.47±0.05 | 0.10±0.04 | -0.25±0.03 | -0.76±0.07 |
|  |  | **M** | 0.91±0.06 | 0.42±0.04 | 0.16±0.04 | -0.17±0.03 | -0.79±0.05 |
| Starting point | **Uncued** | **F** | 0.60±0.02 | 0.55±0.02 | 0.47±0.03 | 0.51±0.04 | 0.35±0.04 |
|  |  | **M** | 0.60±0.01 | 0.53±0.02 | 0.51±0.02 | 0.54±0.02 | 0.43±0.04 |
|  | **Cued** | **F** | 0.63±0.02 | 0.55±0.02 | 0.54±0.02 | 0.51±0.02 | 0.40±0.02 |
|  |  | **M** | 0.57±0.02 | 0.54±0.02 | 0.51±0.01 | 0.50±0.01 | 0.50±0.02 |
| Boundary | **Uncued** | **F** | 3.75±0.14 | 2.67±0.14 | 2.50±0.19 | 2.46±0.25 | 3.01±0.29 |
|  |  | **M** | 3.51±0.09 | 2.55±0.10 | 2.40±0.15 | 2.39±0.11 | 2.87±0.17 |
|  | **Cued** | **F** | 3.60±0.12 | 2.56±0.10 | 2.44±0.07 | 2.64±0.09 | 3.44±0.12 |
|  |  | **M** | 3.16±0.13 | 2.61±0.10 | 2.56±0.08 | 2.94±0.12 | 3.00±0.11 |
| Non-decision time | **Uncued** | **F** | 0.13±0.01 | 0.16±0.01 | 0.16±0.02 | 0.18±0.08 | 0.14±0.04 |
|  |  | **M** | 0.13±0.01 | 0.15±0.02 | 0.23±0.04 | 0.23±0.03 | 0.20±0.02 |
|  | **Cued** | **F** | 0.13±0.01 | 0.14±0.10 | 0.15±0.01 | 0.16±0.02 | 0.13±0.01 |
|  |  | **M** | 0.13±0.01 | 0.19±0.03 | 0.20±0.02 | 0.21±0.03 | 0.20±0.02 |
| szr | **Uncued** | **F** | 0.39±0.03 | 0.40±0.02 | 0.47±0.03 | 0.40±0.03 | 0.38±0.08 |
|  |  | **M** | 0.38±0.02 | 0.43±0.02 | 0.46±0.03 | 0.38±0.02 | 0.37±0.04 |
|  | **Cued** | **F** | 0.36±0.02 | 0.43±0.03 | 0.50±0.03 | 0.47±0.02 | 0.42±0.02 |
|  |  | **M** | 0.36±0.02 | 0.39±0.03 | 0.42±0.02 | 0.38±0.02 | 0.42±0.02 |
| d | **Uncued** | **F** | 0.02±0.006 | 0.03±0.006 | 0.02±0.007 | 0.03±0.009 | 0.04±0.001 |
|  |  | **M** | 0.02±0.003 | 0.04±0.005 | 0.04±0.007 | 0.03±0.009 | 0.02±0.007 |
|  | **Cued** | **F** | 0.03±0.005 | 0.02±0.010 | 0.03±0.006 | 0.02±0.005 | 0.01±0.008 |
|  |  | **M** | 0.03±0.004 | 0.03±0.007 | 0.02±0.005 | 0.02±0.007 | 0.02±0.010 |
